# Targeting an allosteric binding site in the citrate transporter NaCT (SLC13A5)

**DOI:** 10.64898/2026.06.04.730233

**Authors:** Paul Morgan, Wen-An Wang, Giulio Superti-Furga, Avner Schlessinger

## Abstract

The Na^+^-dependent citrate transporter NaCT (SLC13A5) is a key regulator of citrate homeostasis and has emerged as a therapeutic target for metabolic and neurological disease, including the SLC13A5 Epilepsy, a rare disease marked by severe sezures and neurodevelopmental delays. Current NaCT inhibitors are substrate-like molecules that competitively bind the substrate binding site. In this study, we identify previously unknown small molecule inhibitors of NaCT by targeting a putative allosteric site located at the dimer interface. We performed a virtual screen of 3.5 million compounds from the ZINC20 database against this site and selected 54 candidates for experimental testing using a cell-based citrate uptake assay. Through initial experiments, we identified three weak inhibitors, and subsequent evaluation of 26 structurally related analogs yielded six compounds with improved potency (IC_50_ = 12.78 μM and 15.49 μM). We then performed further analysis of the putative binding site by integrating structural data with deep mutational scanning evidence and comparisons with homolog structures. This analysis highlighted the importance of key residues (e.g., Phe362) in ligand modulation. These findings reveal a promising allosteric pocket and establish a chemically distinct series of NaCT inhibitors, providing a foundation for rational development of pharmacological modulators of NaCT function.

## INTRODUCTION

The sodium-dependent citrate transporter SLC13A5 (NaCT) transports intermediates of the tricarboxylic acid (TCA) cycle, including citrate and succinate, across the plasma membrane, thereby regulating nutrient availability for metabolic and neurophysiological processes.^1-6^ NaCT is highly expressed in the liver and brain, where it supplies citrate to biosynthetic pathways and modulates enzymes such as acetyl-CoA carboxylase and phosphofructokinase-1.^1,2^ In metabolic tissues, reduced NaCT activity has been associated with enhanced mitochondrial biogenesis, decreased lipogenesis, and protection from diet- or age-induced metabolic dysfunction, supporting therapeutic strategies based on NaCT inhibition.^7^

In contrast, loss-of-function mutations in SLC13A5 cause SLC13A5 Epilepsy, a neonatal-onset epileptic encephalopathy characterized by severe neurodevelopmental impairment and elevated systemic citrate levels.^5,6^ More than 50 pathogenic variants have been identified, many of which disrupt transporter function through impaired folding, trafficking, or conformational dynamics.^8-11^ Structural and computational studies, including cryo-EM analyses, have begun to elucidate how these mutations perturb the transport cycle, providing a framework for therapeutic intervention.^3,5,8,11-13^

Together, these opposing disease contexts highlight a central challenge: NaCT function must be selectively modulated rather than uniformly inhibited, depending on the pathological setting. This underscores the need for pharmacological strategies capable of both suppressing and restoring transporter activity.

To date, most small-molecule efforts targeting NaCT have largely focused on the orthosteric substrate-binding site. Early inhibitors identified through homology modeling and docking exhibited limited potency and selectivity.^3,14^ More advanced compounds, including Pfizer’s dicarboxylate series (PF-06649298/PF2 and PF-06761281), bind within the canonical citrate-binding pocket, engaging conserved SNT motifs and stabilizing an inward-facing conformation.^5,16^ Although potent, these compounds are structurally similar to citrate and often display substrate-like behavior, functioning as both inhibitors and low-affinity transported substrates.^15^ More recently, non-acidic NaCT inhibitors, including BI01383298 (PMID: 42049193) and ETG-5773 (PMID: 36005604), have provided more drug-like starting points for future therapeutic development.

Cryo-EM structures confirm that PF2 occupies the substrate-binding site, coordinating Na^+^ ions and residues that govern selectivity within the SLC13 family.^15^ Additional NaCT ligands reported to date, including those identified through high-throughput screening or rational design, similarly target this orthosteric site, leaving alternative binding pockets largely unexplored. ^15^ By contrast, studies of related transporters have demonstrated that allosteric sites can support diverse modes of modulation, including inhibition, stabilization, and activation, highlighting the potential of non-orthosteric approaches for NaCT.^1,2^

Here, we report the discovery of a chemically distinct class of NaCT inhibitors identified through structure-based virtual screening of 3.5 million compounds from ZINC20 targeting a putative allosteric cavity. Through a two-stage experimental validation using a cell-based citrate uptake assay, we identified novel inhibitors predicted to bind at the dimer interface, a region spatially distinct from the canonical substrate-binding site. These findings establish an alternative modality for NaCT modulation and provide a framework for the development of both inhibitory and function-restoring ligands.

## RESULTS

### Identification of a putative allosteric pocket at the NaCT dimer interface

To predict previously unknown binding sites on the surface of NaCT we used the binding site prediction methods CAVIAR^16^ and SiteMap^17^ with the inward-facing cryo-EM structure as input.^2^ Both methods consistently revealed a well-defined pocket at the dimer interface, positioned between the transport and scaffold domains of each protomer (Protomer A and Protomer B; Fig. 1). The cavity exhibited favorable physicochemical features for ligand engagement, including size (∼334 Å^3^), depth, and hydrophobicity (54% hydrophobic; ligandability score = 1), consistent with a druggable allosteric site that could modulate transporter function by stabilizing or destabilizing dimer interactions depending on the bound ligand. These observations suggested that binding at this region could influence dimer stability or the conformational transitions required for an elevator-type transport.^18-20^ The interfacial cavity consists of two main subpockets, separated by a narrow channel with sufficient width to accommodate a core linker. Notably, structural comparison with related SLC13 family members (NaDC1 and NaS1) revealed that the cavity corresponds to a region previously implicated in lipid regulation within those homologs, further supporting its potential functional importance.^1^

**Fig. 1.**
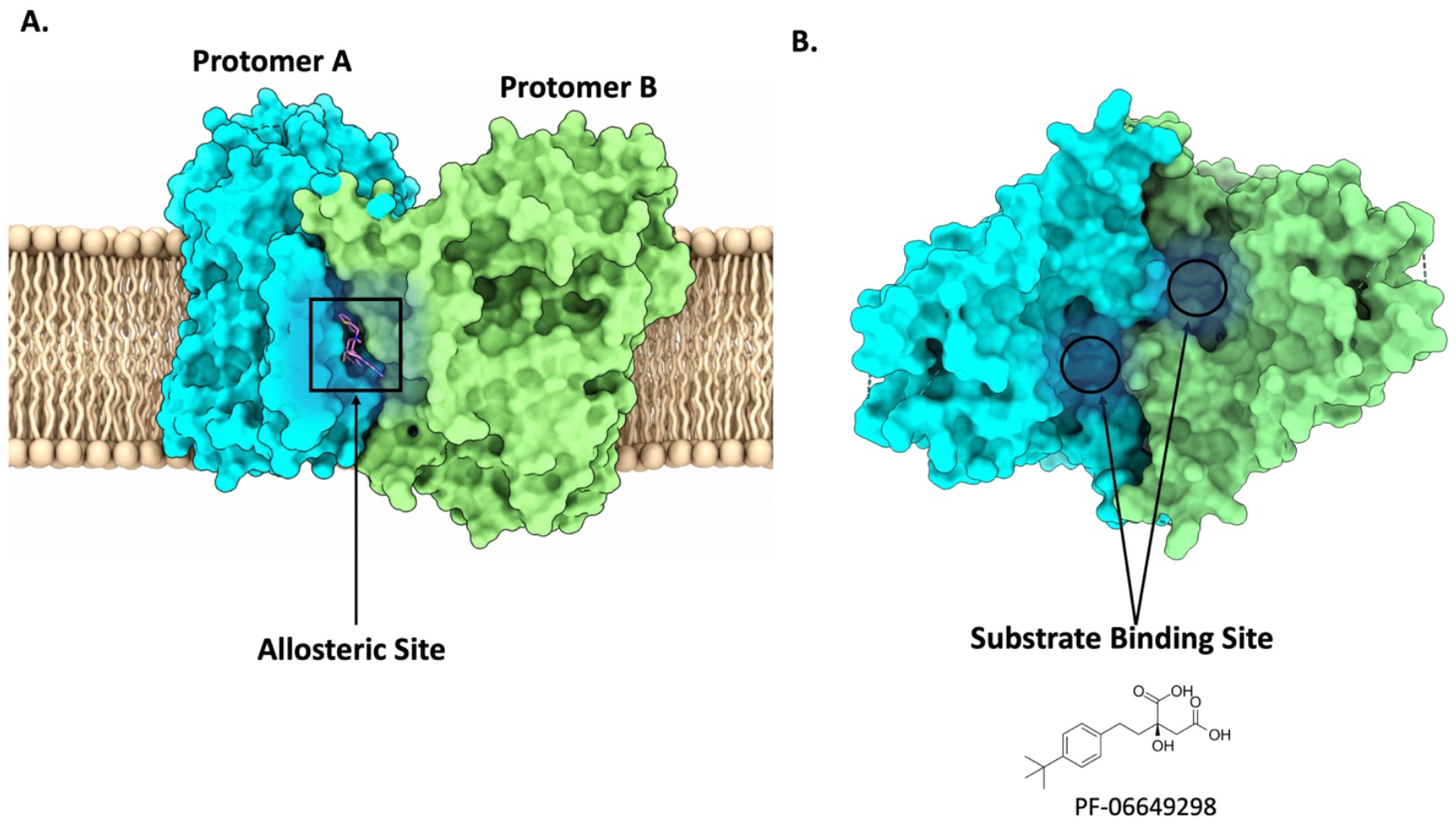
Predicted allosteric site. (A) Surface representation of the human NaCT(SLC13A5) dimer in the inward-facing conformation derived from cryo-EM structure (PDB 7JSJ, 3.1 Å resolution). Protomer A and Protomer B are shown are shown in cyan and green, respectively. The predicted allosteric binding site, highlighted by the boxed region, is located at the interface between the two protomers within the membrane environment. The bound ligand is shown as magenta sticks. (B) Top-down view of the NaCT dimer illustrating the central substrate-binding cavities within each protomer (black circles). The interfacial allosteric site is spatially distinct from the canonical substrate-binding site.

### Virtual screening identifies initial hit compounds

We used Glide to virtually screen 3.5 million lead-like ZINC20 molecules against the putative allosteric site.^21^ After clustering the top 2,000 scoring ligands by chemical similarity, we used a structure-guided selection strategy that prioritized ligands capable of engaging both protomers within the interfacial binding site which could potentially have a stabilizing or destabilizing effect. We selected molecules predicted to form π–π stacking interactions with F362 in protomer A (F_A_362) and/or F18 in protomer B (F_B_18), complemented by polar contacts with the side chains of Y48_B_ and/or Y_A_392 (Fig. 2). These interactions were hypothesized to cooperatively reinforce inter-protomer coupling cooperatively, while the largely rigid diaspiro sub-scaffold was intentionally positioned deep within the hydrophobic core of the pocket to serve as a conformational anchor. 30 structurally diverse representatives were purchased for experimental testing. These compounds were evaluated using a citrate biosensor assay capable of detecting NaCT-dependent citrate uptake.^4^ Three compounds reproducibly inhibited citrate transport in the low-to-mid micromolar range. In the top docking pose of compound 20 (IC_50_ = 25.37 µM), the diaspiro core is buried deep within the pocket, with the pyridine and thiophene substituents forming parallel and T-shaped π–π interactions with F362_A_ and F18_B_, respectively (Fig. 2).

**Fig. 2:**
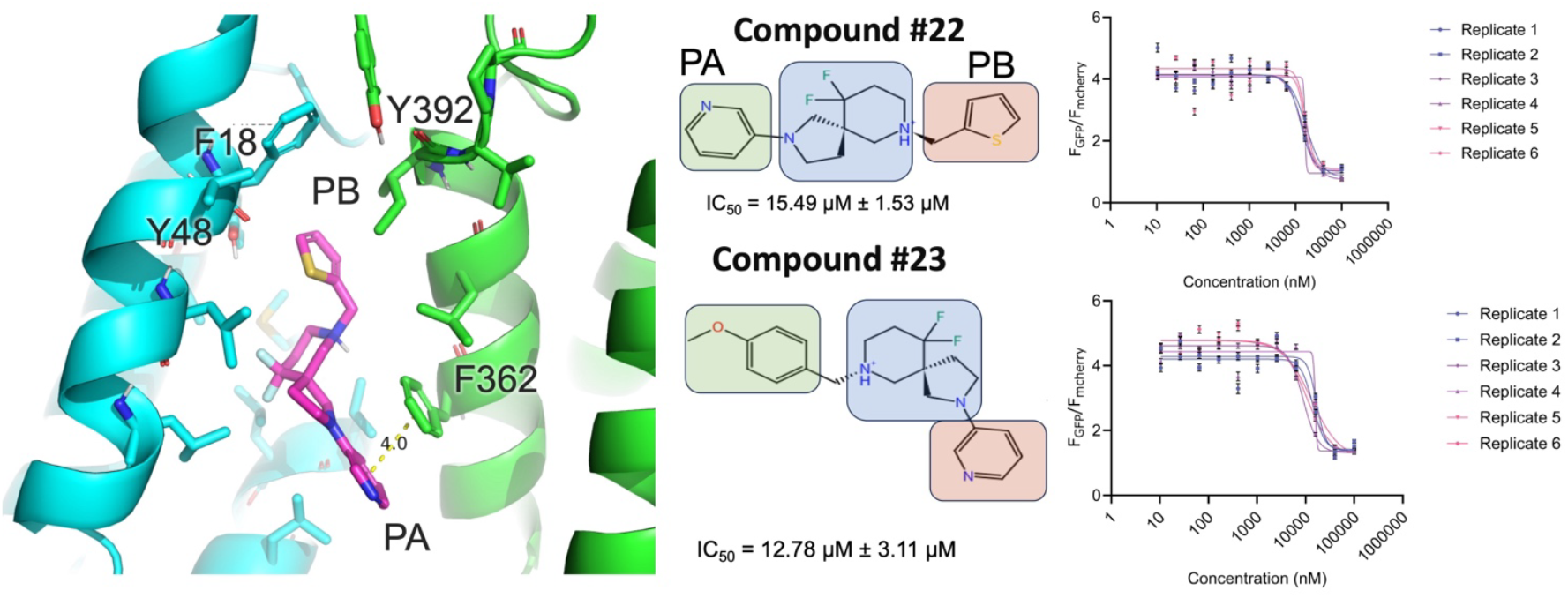
Small-molecule hits targeting the NaCT dimer interface. (**A**) Predicted binding mode of Compound 22. Shown is a sideview of the NaCT dimer interface in the inward-facing conformation structure where the subunits are in green and cyan ribbons; key binding site residues are shown as sticks. Compound 2 (in magenta) is predicted to engage both monomers by occupying two adjacent subpockets (PA and PB); hydrogen bonds are represented with yellow dashes. Phe362 as a key determinant of inhibitor sensitivity, consistent with its predicted π-stacking interactions with PB-directed aromatic groups (**B**) 2D structures of the hit compounds with key moieties highlighted in green, blue, and yellow. A central diazaspiro moiety appears critical for binding (blue), functioning both as a structural anchor within the interfacial cavity and as the linker that positions the PA and PB interacting groups for simultaneous engagement. (**C**) Experimental testing of top hits. IC_50_ curves were generated using the fluorescence-based citrate biosensor assay, and no cytotoxicity was observed at relevant concentrations. Dose-response analysis of compounds #22 and #23. Data represent mean ± SEM (n = 6 independent experiments). Curves show nonlinear regression fits. (four-parameter logistic model). IC_50_ values are reported as mean ± SEM.

We used Glide to virtually screen 3.5 million lead-like ZINC20 molecules against the putative allosteric site. From this screen, three compounds reproducibly inhibited NaCT-mediated citrate uptake in the low-to-mid micromolar range, with the top hit (compound 20) exhibiting an IC_50_ of 25.37 µM. ^21^ After clustering the top 2,000 scoring ligands by chemical similarity, we applied a structure-guided selection strategy that prioritized ligands capable of engaging both protomers within the interfacial binding site, with the goal of modulating inter-protomer coupling. Selected candidates were predicted to form π–π stacking interactions with F362 in protomer A (FA362) and/or F18 in protomer B (FB18), complemented by polar contacts with Y48B and/or YA392 (Fig. 2). In the top docking pose of compound 20, the diaspiro core is buried deep within the hydrophobic pocket, while the pyridine and thiophene substituents form parallel and T-shaped π–π interactions with F362A and F18B, respectively (Fig. 2). This binding mode positions the rigid diaspiro scaffold as a conformational anchor within the interfacial cavity. A total of 30 structurally diverse compounds were purchased and evaluated using a citrate biosensor assay to measure NaCT-dependent citrate uptake. ^4^

### Structure-Activity-Relationship (SAR) by catalog

To refine the initial hits, we purchased and tested 26 analogs of the most promising scaffold (compound 22; Fig. 2).^22^ Six analogs demonstrated substantially improved inhibition, including two with IC_50_ values of 12.78 μM and 15.49 μM, marking an improvement over the initial screening hits. Importantly, none of the compounds exhibited cytotoxicity in viability assays, supporting their suitability for further optimization. Clear SAR trends emerged from this series: hydrophobic expansion within subpocket PB consistently improved activity, aligning with the pocket’s largely nonpolar character, while electron-rich aromatic groups positioned toward PA enhanced potency by forming favorable π-stacking interactions with Phe362. Docking analyses revealed that the majority of active molecules adopted conformations bridging PA and PB, with structural features consistent with allosteric modulation at the dimer interface.^23^

### Mutational evidence supports functional relevance of the pocket

To probe the mechanistic role of the interfacial cavity, we analyzed the predicted binding site and mode of interaction of inhibitors in the context of a recent comprehensive analysis that used deep mutational scanning (DMS) to quantify the functional consequences of all possible missense variants in SLC13A5.^24^ Our data suggests that mutation of Phe362, a residue predicted to form π–π interactions with the PA-facing aromatic moiety of active molecules, significantly altered inhibitor sensitivity. This further supports a direct functional role of this pocket in small-molecule modulation.

Together, these results establish the interfacial region of NaCT as a promising allosteric site and the diazaspiro scaffold a unique class of small-molecule inhibitors capable of modulating NaCT activity through inter-protomer interactions.

## DISCUSSION

Targeting the sodium-coupled citrate transporter SLC13A5 (NaCT) presents a unique therapeutic challenge due to its context-dependent role in human disease.^1,13,15,22,25^ In metabolic disorders such as hyperlipidemia and fatty liver disease, reduced NaCT activity has been associated with beneficial effects on lipid metabolism, motivating the development of inhibitors. In contrast, loss-of-function mutations in SLC13A5 cause SLC13A5 Epilepsy, where restoration or enhancement of transporter function represents a therapeutic goal. These opposing disease contexts highlight the need for pharmacological strategies capable of bidirectional modulation, rather than uniform inhibition.

To date, efforts to target NaCT have largely relied on substrate-like compounds that bind the orthosteric citrate-binding site.^18^ While some of these compounds have demonstrated clinical promise, their structural similarity to citrate and reliance on conserved binding motifs have limited selectivity and constrained chemical diversity. More importantly, orthosteric strategies inherently favor inhibition and provide limited opportunities to stabilize or enhance transporter function.^18^

These limitations motivate the exploration of alternative binding sites and modalities. In this study, we targeted a previously unexplored interfacial pocket at the NaCT dimer interface, hypothesizing that ligands engaging this region could modulate inter-protomer interactions and thereby influence transporter function through non-orthosteric mechanisms. Such sites may, in principle, support both inhibitory and activating modes of action, depending on how ligand binding alters conformational equilibria.

Consistent with this framework, we identified a chemically distinct series of NaCT inhibitors that expand the accessible chemical space beyond di- and tri-carboxylate-derived scaffolds. A defining feature of these compounds is their bipartite architecture, in which linked aromatic and heterocyclic moieties span two adjacent subpockets (Fig. 2). Preliminary SAR indicates that both regions contribute to activity, consistent with a mechanism in which ligands act as molecular clamps or wedges that alter inter-protomer coupling. While the precise binding interactions remain to be experimentally validated, inhibitor activity aligns with the modulatory role of Phe362 identified through deep mutational scanning.^24^ Together, these observations support a shared, non-orthosteric mechanism of action.

Importantly, the identification of inhibitors at this interfacial site provides a conceptual foundation for broader modulation strategies. Small-molecule correctors of CFTR in cystic fibrosis demonstrate that transporter function can be restored by ligands that stabilize specific conformational states rather than directly competing with substrate binding.^26^ Similar approaches may be applicable to SLC transporters, including SLC13A5, for which many disease-associated variants retain partial transport competence but exhibit defects in folding, trafficking, or conformational stability.^9,27,28^ In this context, ligands targeting the dimer interface could act not only as inhibitors that disrupt protomer coupling, but also as “molecular glues” that stabilize productive conformations and restore function. Although the present study identified inhibitors, related scaffolds could potentially be optimized as pharmacochaperones for SLC13A5 Deficiency Disorder.

Despite these advances, several limitations remain. Structural validation of the proposed binding site will be critical, and future studies employing ligand-bound cryo-EM, HDX-MS, or cross-linking approaches will be necessary to define the molecular interactions underlying activity. Selectivity profiling across the SLC13 family will also be required to determine whether the dimer interface provides a viable route to subtype-selective modulation.^29^ Furthermore, while initial SAR yielded low-micromolar inhibitors, further medicinal chemistry efforts aimed at optimizing pocket engagement, enhancing π-stacking interactions, and improving physicochemical properties will be necessary to achieve higher potency. Finally, systematic evaluation of these compounds across pathogenic SLC13A5 variants will be essential to assess their potential as stabilizers or folding correctors.

In summary, this study establishes an alternative strategy for targeting NaCT by demonstrating that an allosteric pocket at the dimer interface can be exploited to achieve non-orthosteric modulation. By introducing a chemically novel scaffold and a new binding site, these findings expand the pharmacological landscape of NaCT and lay the groundwork for the development of both inhibitory and function-restoring ligands for metabolic disorders and SLC13A5 Deficiency Disorder.

## Methods

### Binding site identification

Potential ligand binding sites were identified using CAVIAR (default parameters) and SiteMap (Schrödinger).^17^ Both methods predicted eight candidate cavities, which were evaluated based on size, enclosure, and hydrophobic character. Among these, a pocket located at the SLC13A5 dimeric interface ranked highly in both analyses and exhibited favorable druggability features. The site spans the interprotomer interface positioning it to modulate dimeric coupling and conformational dynamics. With a volume of ∼334 Å^3^ and 54% hydrophobicity, the cavity provides an environment conducive to high-affinity ligand binding.^16,17^ Its favorable ligandability score further prioritizes this site as a structurally and functionally relevant allosteric pocket for therapeutic targeting.^18^ Given its consistent identification across methods and its location at the dimer interface, this site was selected for subsequent docking and virtual screening.

### Molecular docking

Molecular docking was performed using the Glide module in the Schrödinger Suite.^21^ All ligands were docked to the inward-facing cryo-EM structure of NaCT (PDB ID: 7JSJ).^30^ The protein structure was prepared with the Protein Preparation Wizard in Maestro using default settings. The ligand-binding site was defined according to cavity coordinates identified by CAVIAR, an automated cavity detection algorithm, and a receptor grid was subsequently generated using the Maestro Receptor Grid Generation tool. A total of 3.5 million compounds from the ZINC20 lead-like library were prepared for docking using LigPrep (Schrödinger).^21^ The top 2,000 scoring compounds were clustered based on chemical similarity, and docking poses and interactions were analyzed using PyMOL.^22^ Top-scoring compounds were clustered using ECFP4 fingerprints^31^ and Tanimoto similarity (0.8 cutoff),^32^ and represented molecules were selected to maximize chemical diversity.^33^ Visual inspection prioritized ligands exhibiting consistent binding modes, favorable shape complementarity, and inter-protomer interactions. Docking poses were specifically evaluated for interactions with key interfacial residues F362 and F18, as well as polar contacts involving Y392 and Y48. Compounds were further enriched for aromatic substituents engaging predicted π–π interactions at the interfacial pocket and for rigid core scaffolds deeply buried within the hydrophobic region, consistent with a conformational anchoring role. From these clusters, 30 chemically diverse compounds were selected for experimental validation using a fluorescence-based citrate biosensor assay.

### Citrate biosensor assay

The citrate biosensor assay was performed as previously described.^24^ Briefly, the Jump In T-Rex HEK293 (Thermo Fisher Scientific, no. A15008) cells stably overexpressing SLC13A5, were seeded on poly-lysine–coated 96-well plates (PerkinElmer, 6005550), at 10,000 cells per well. After 24 hours in culture, the cells were transfected with pCMV-Citron1-p2a-mCherry for 24 hours using Lipofectamine 3000 Transfection Reagent (Thermo Fisher Scientific, no. L3000001) according to the manufacturer’s protocol. Cells were then treated with or without doxycycline (2 mg/ml) for inducing SLC13A5 expression. At this point, cells were also treated with different compounds at different concentrations and imaging experiments were performed after 24 hours.

On the day of imaging experiments, culture medium was exchanged for imaging buffer (Hanks’ balanced salt solution (Thermo Fisher Scientific, no. 14025092) with 25 mM Hepes (Gibco, 15630056)) containing varying concentrations of citrate and corresponding compounds, and was incubated for 10 min at 37°C with 5% CO_2_ and 95% humidity. Images were captured with six fields per well at ×40 magnification on the PerkinElmer Opera high-throughput microscope using the channels for green fluorescent protein (GFP; λex, 488 nm/λem, 500 to 550 nm) to measure Citron1 and mCherry (λex, 561 nm/λem, 570 to 630 nm). Quantification of Citron1 and mCherry fluorescence were performed with ImageJ using JIPipe plugin.^34^ The region of interest for each cell was defined using the mCherry fluorescence, and the average fluorescence intensity for both mCherry and GFP were measured within those regions. The ratio, calculated based on *F*_GFP_/*F*_mCherry_, was used as a measure of citrate in the cell and the ratio over citrate concentration was used to calculate an apparent *K*_m_, or IC_50_.

### Citrate toxicity assay

The citrate toxicity assay was performed as previously.^24^ Briefly cells were seeded on poly-lysine– coated 96-well plates (PerkinElmer, 6005550), at 10,000 cells per well and treated with or without doxycycline (2 mg/ml) for inducing *SLC13A5* expression. After 24 hours, we incubated the cells with varying concentrations of compounds for 48 hours and performed a CellTiter-Glo Luminescent Cell Viability Assay (Promega, G7570), to test for cell survival, according to the manufacturer’s protocol.

## References

1. Chi X, Chen Y, Li Y, et al. Cryo-EM structures of the human NaS1 and NaDC1 transporters revealed the elevator transport and allosteric regulation mechanism. Science Advances. 2024;10(13):eadl3685. doi:doi:10.1126/sciadv.adl3685

2. Sauer DB, Song J, Wang B, et al. Structure and inhibition mechanism of the human citrate transporter NaCT. Nature. 2021/03/01 2021;591(7848):157–161. doi:10.1038/s41586-021-03230-x

3. Kumar A, Cordes T, Thalacker-Mercer AE, Pajor AM, Murphy AN, Metallo CM. NaCT/SLC13A5 facilitates citrate import and metabolism under nutrient-limited conditions. Cell Rep. Sep 14 2021;36(11):109701. doi:10.1016/j.celrep.2021.109701

4. Zhao Y, Shen Y, Wen Y, Campbell RE. High-Performance Intensiometric Direct-and Inverse-Response Genetically Encoded Biosensors for Citrate. ACS Central Science. 2020/08/26 2020;6(8):1441–1450. doi:10.1021/acscentsci.0c00518

5. Klotz J, Porter BE, Colas C, Schlessinger A, Pajor AM. Mutations in the Na(+)/citrate cotransporter NaCT (SLC13A5) in pediatric patients with epilepsy and developmental delay. Mol Med. May 26 2016;22:310–21. doi:10.2119/molmed.2016.00077

6. Thevenon J, Milh M, Feillet F, et al. Mutations in SLC13A5 cause autosomal-recessive epileptic encephalopathy with seizure onset in the first days of life. Am J Hum Genet. Jul 3 2014;95(1):113–20. doi:10.1016/j.ajhg.2014.06.006

7. Birkenfeld AL, Lee HY, Guebre-Egziabher F, et al. Deletion of the mammalian INDY homolog mimics aspects of dietary restriction and protects against adiposity and insulin resistance in mice. Cell Metab. Aug 3 2011;14(2):184–95. doi:10.1016/j.cmet.2011.06.009

8. Schlessinger A, Zatorski N, Hutchinson K, Colas C. Targeting SLC transporters: small molecules as modulators and therapeutic opportunities. Trends Biochem Sci. Sep 2023;48(9):801–814. doi:10.1016/j.tibs.2023.05.011

9. Colas C, Ung PM, Schlessinger A. SLC Transporters: Structure, Function, and Drug Discovery. Medchemcomm. Jun 1 2016;7(6):1069–1081. doi:10.1039/c6md00005c

10. Schlessinger A, Sun NN, Colas C, Pajor AM. Determinants of substrate and cation transport in the human Na+/dicarboxylate cotransporter NaDC3. J Biol Chem. Jun 13 2014;289(24):16998–7008. doi:10.1074/jbc.M114.554790

11. Hardies K, de Kovel CG, Weckhuysen S, et al. Recessive mutations in SLC13A5 result in a loss of citrate transport and cause neonatal epilepsy, developmental delay and teeth hypoplasia. Brain. Nov 2015;138(Pt 11):3238–50. doi:10.1093/brain/awv263

12. Schlessinger A, Welch MA, van Vlijmen H, Korzekwa K, Swaan PW, Matsson P. Molecular Modeling of Drug-Transporter Interactions-An International Transporter Consortium Perspective. Clin Pharmacol Ther. Nov 2018;104(5):818–835. doi:10.1002/cpt.1174

13. Pajor AM, de Oliveira CA, Song K, Huard K, Shanmugasundaram V, Erion DM. Molecular Basis for Inhibition of the Na+/Citrate Transporter NaCT (SLC13A5) by Dicarboxylate Inhibitors. Mol Pharmacol. Dec 2016;90(6):755–765. doi:10.1124/mol.116.105049

14. Colas C, Pajor AM, Schlessinger A. Structure-Based Identification of Inhibitors for the SLC13 Family of Na(+)/Dicarboxylate Cotransporters. Biochemistry. Aug 11 2015;54(31):4900–8. doi:10.1021/acs.biochem.5b00388

15. Huard K, Brown J, Jones JC, et al. Discovery and characterization of novel inhibitors of the sodium-coupled citrate transporter (NaCT or SLC13A5). Scientific Reports. 2015/12/01 2015;5(1):17391. doi:10.1038/srep17391

16. Marchand JR, Pirard B, Ertl P, Sirockin F. CAVIAR: a method for automatic cavity detection, description and decomposition into subcavities. J Comput Aided Mol Des. Jun 2021;35(6):737–750. doi:10.1007/s10822-021-00390-w

17. Halgren TA. Identifying and characterizing binding sites and assessing druggability. J Chem Inf Model. Feb 2009;49(2):377–89. doi:10.1021/ci800324m

18. Kinz-Thompson CD, Lopez-Redondo ML, Mulligan C, et al. Elevator mechanism dynamics in a sodium-coupled dicarboxylate transporter. bioRxiv. 2025:2022.05.01.490196. doi:10.1101/2022.05.01.490196

19. Mulligan C, Fenollar-Ferrer C, Fitzgerald GA, et al. The bacterial dicarboxylate transporter VcINDY uses a two-domain elevator-type mechanism. Nature Structural & Molecular Biology. 2016/03/01 2016;23(3):256–263. doi:10.1038/nsmb.3166

20. Vergara-Jaque A, Fenollar-Ferrer C, Mulligan C, Mindell JA, Forrest LR. Family resemblances: A common fold for some dimeric ion-coupled secondary transporters. Journal of General Physiology. 2015;146(5):423–434. doi:10.1085/jgp.201511481

21. Friesner RA, Murphy RB, Repasky MP, et al. Extra Precision Glide: Docking and Scoring Incorporating a Model of Hydrophobic Enclosure for Protein−Ligand Complexes. Journal of Medicinal Chemistry. 2006/10/01 2006;49(21):6177–6196. doi:10.1021/jm051256o

22. Irwin JJ, Tang KG, Young J, et al. ZINC20—A Free Ultralarge-Scale Chemical Database for Ligand Discovery. Journal of Chemical Information and Modeling. 2020/12/28 2020;60(12):6065–6073. doi:10.1021/acs.jcim.0c00675

23. Qiu Z, Wang W, Nie Y, et al. Allosteric modulation of dimeric GPR3 by ligands in the dimerization interface. eLife Sciences Publications, Ltd; 2025.

24. Wang W-A, Ferrada E, Klimek C, et al. Large-scale experimental assessment of variant effects on the structure and function of the citrate transporter SLC13A5. Science Advances. 2025;11(26):eadx3011. doi:doi:10.1126/sciadv.adx3011

25. Huard K, Gosset JR, Montgomery JI, et al. Optimization of a Dicarboxylic Series for in Vivo Inhibition of Citrate Transport by the Solute Carrier 13 (SLC13) Family. Journal of Medicinal Chemistry. 2016/02/11 2016;59(3):1165–1175. doi:10.1021/acs.jmedchem.5b01752

26. Levring J, Terry DS, Kilic Z, Fitzgerald G, Blanchard SC, Chen J. CFTR function, pathology and pharmacology at single-molecule resolution. Nature. 2023/04/01 2023;616(7957):606–614. doi:10.1038/s41586-023-05854-7

27. Zhang Y, Zhang Y, Sun K, Meng Z, Chen L. The SLC transporter in nutrient and metabolic sensing, regulation, and drug development. J Mol Cell Biol. Jan 1 2019;11(1):1–13. doi:10.1093/jmcb/mjy052

28. Schlessinger A, Khuri N, Giacomini KM, Sali A. Molecular modeling and ligand docking for solute carrier (SLC) transporters. Curr Top Med Chem. 2013;13(7):843–56. doi:10.2174/1568026611313070007

29. Colas C, Pajor AM, Schlessinger A. Structural Characterization of Substrate Transport Selectivity of the SLC13 Family of Na^+^/Dicarboxylate Cotransporters. Biophysical Journal. 2016;110(3):627a. doi:10.1016/j.bpj.2015.11.3362

30. Halgren TA, Murphy RB, Friesner RA, et al. Glide: A New Approach for Rapid, Accurate Docking and Scoring. 2. Enrichment Factors in Database Screening. Journal of Medicinal Chemistry. 2004/03/01 2004;47(7):1750–1759. doi:10.1021/jm030644s

31. Rogers D, Hahn M. Extended-Connectivity Fingerprints. Journal of Chemical Information and Modeling. 2010/05/24 2010;50(5):742–754. doi:10.1021/ci100050t

32. Schottlender G, Prieto JM, Marti MA, Fernández Do Porto D. Beyond Tanimoto: a learned bioactivity similarity index enhances ligand discovery. Front Bioinform. 2025;5:1695353. doi:10.3389/fbinf.2025.1695353

33. Talevi A, Bellera CL. Clustering of small molecules: new perspectives and their impact on natural product lead discovery. Perspective. Frontiers in Natural Products. 2024-March-15 2024; Volume 3-2024 doi:10.3389/fntpr.2024.1367537

34. Gerst R, Cseresnyés Z, Figge MT. JIPipe: visual batch processing for ImageJ. Nature Methods. 2023/02/01 2023;20(2):168–169. doi:10.1038/s41592-022-01744-4

